# Regulation of the *ACE2* locus in human airways cells

**DOI:** 10.1101/2020.10.04.325415

**Authors:** Hye Kyung Lee, Olive Jung, Lothar Hennighausen

**Affiliations:** Laboratory of Genetics and Physiology, National Institute of Diabetes, Digestive and Kidney Diseases, National Institutes of Health, Bethesda, Maryland 20892; Division of Preclinical Innovation, National Center for Advancing Translational Sciences, National Institutes of Health, Bethesda, Maryland 20892

## Abstract

The angiotensin-converting enzyme 2 (ACE2) receptor is the gateway for SARS-CoV-2 to airway epithelium^1,2^ and the strong inflammatory response after viral infection is a hallmark in COVID-19 patients. Deciphering the regulation of the ACE2 gene is paramount for understanding the cell tropism of SARS-CoV-2 infection. Here we identify candidate regulatory elements in the *ACE2* locus in human primary airway cells and lung tissue. Activating histone and promoter marks and Pol II loading characterize the intronic *dACE2* and define novel candidate enhancers distal to the genuine *ACE2* promoter and within additional introns. *dACE2*, and to a lesser extent *ACE2*, RNA levels increased in primary bronchial cells treated with interferons and this induction was mitigated by Janus kinase (JAK) inhibitors that are used therapeutically in COVID-19 patients. Our analyses provide insight into regulatory elements governing the *ACE2* locus and highlight that JAK inhibitors are suitable tools to suppress interferon-activated genetic programs in bronchial cells.

## Introduction

Recent studies^3,4^ have identified a novel short form of ACE2, called dACE2, that originates from an intronic promoter activated by interferons. Onabajo et al.^4^ used ENCODE data for chromatin modification marks (H3K4me3, H3K4me1 and H3K27ac) as well as DNase I hypersensitive (DHS) sites in cell lines to label putative regulatory elements at the newly identified exon (ex1c) located within intron 9 of the *ACE2* gene. However, no regulatory elements were detected in the vicinity of the 5’ end of the full-length transcript encoding biologically active ACE2, and in sequences distal to the genuine promoter. Since these data sets were obtained from a wide range of cell lines and not from human primary airway cells, the principal target of SARS-CoV-2, they might not present a comprehensive picture of the regulatory regions controlling expression of the *dACE2* and the full-length *ACE2* transcripts in bronchial tissue.

## Results

To comprehensively identify the genetic elements controlling the extended *ACE2* locus, with an emphasis on its interferon response, we focused on human primary Small Airway Epithelial Cells (SAEC), which express a wide range of cytokine receptors and key mechanistic components of the executing JAK/STAT signal transduction pathway (Supplementary Table 1). We stimulated SAECs with interferon type I (IFN α and IFNβ), type II (IFNγ) and type III (IFNλ) as well as with growth hormone (GH), Interleukin 6 (IL6) and IL7, followed by RNA-seq transcriptome analyses (Supplementary Tables 2-8). Increased *ACE2* expression was obtained with the interferons but not with GH, IL6 and IL7 (Supplementary Figure 1). However, the induction was less than that seen for classical interferon stimulated genes (ISG), such as *STAT1*. In agreement with earlier studies^3,4^, we detected the novel *dACE2* N-terminal exon (ex1c) within intron 9 of the *ACE2* gene (Supplementary Figure 1). To obtain more definitive information on the interferon response of the *dACE2* and *ACE2* promoters, we used RNA-seq and determined the respective read counts over the three alternative first exons (Figure 1a and Supplementary Figure 1d). While the increase of RNA-seq reads induced by IFN α/β was highest (∼25-fold) over ex1c, a lesser, yet significant, ∼2-10-fold increase was detected over ex1a and ex1b, supporting the notion that expression of the full-length *ACE2* transcript is also under interferon control. As an independent assay we used qRT-PCR and determined that IFN α/β stimulation led to a 8 to 15-fold increase of *dACE2* and an approximately ∼3-fold increase of ACE2 RNA (Figure 1b). However, the degree of induction of either form was lower than that seen for *bona fide* ISGs (Supplementary Figure 1). Previous studies in normal human bronchial epithelium (NHBE) did not reveal an interferon response of the native *ACE2* promoter^3,4^ suggesting differences between cell types or growth conditions. The mouse *Ace2* gene is induced by cytokines through a STAT5-based enhancer in the second intron^5^ and a DHS site is located in the equivalent location in the human *ACE2* gene in SAEC and lung tissue. This suggests the presence of additional regulatory elements controlling expression of the full-length *ACE2* mRNA.

**Figure 1.**
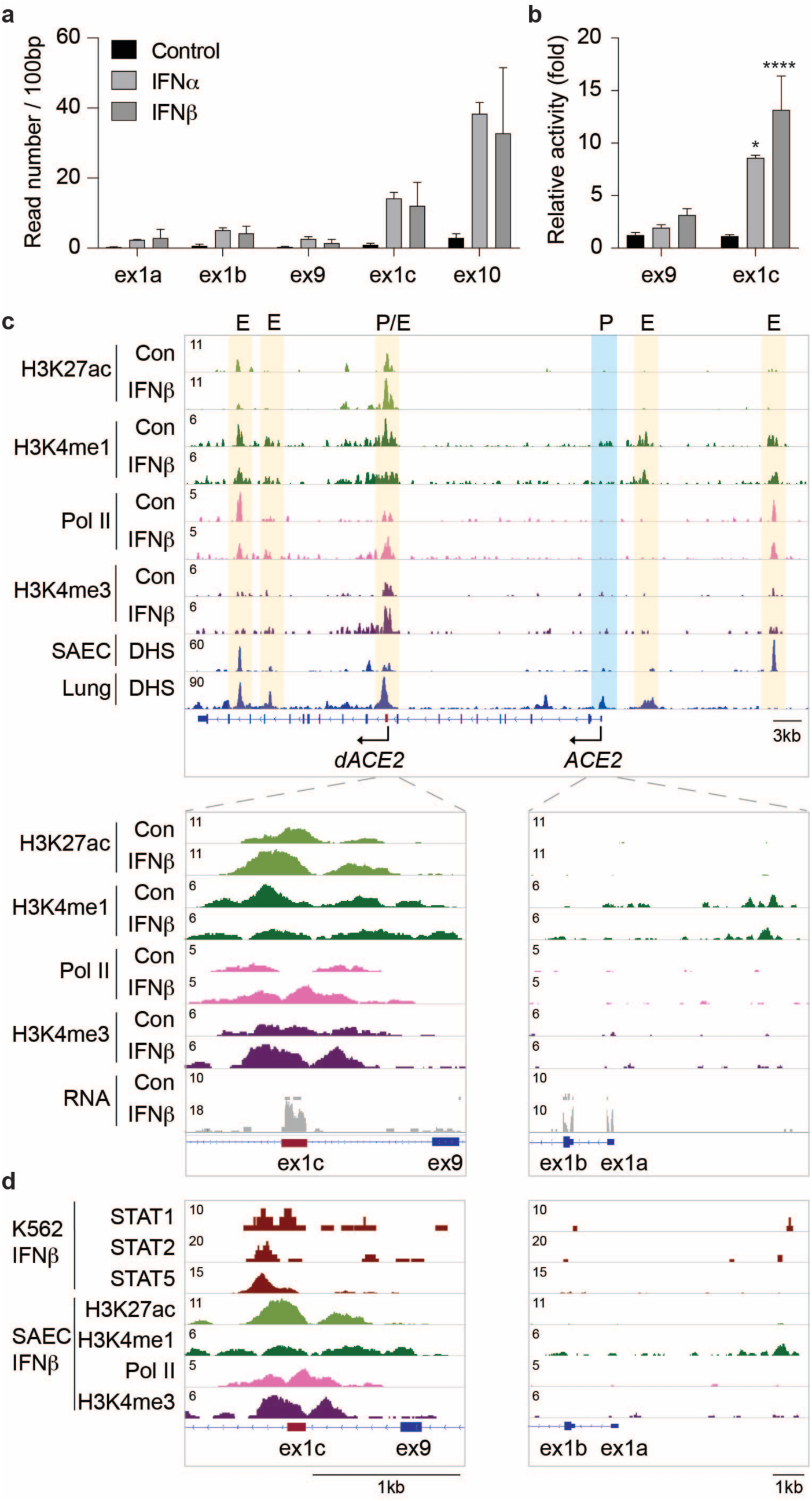
Regulation of the *ACE2* gene in primary airway epithelium. **a**. SAECs were cultured in the absence and presence of interferon alpha (IFN α) and beta (IFNβ) followed by RNA-seq analyses. The reads covering key exons (1a, 1b, 9, 1c and 10) are shown. **b**. mRNA levels of exon9 and exon1c were measured using qRT-PCR. Results are shown as the means ± s.e.m. of independent biological replicates (Control and IFNβ, *n* = 9; IFN α, *n* = 3). Two-way ANOVA with followed by Tukey’s multiple comparisons test was used to evaluate the statistical significance of differences. **c**. ChIP-seq experiments for the histone marks H3K4me3 (promoter), H3K4me1 (enhancers), H3K27ac (active genes) and Pol II loading. The DHS data were obtained from ENCODE^6,7^. Yellow shade, candidate enhancers and blue shade, predicted promoter. The P/E region within intron 9 probably constitutes a combined promoter/enhancer. **d**. Putative STAT5 enhancer in the *ACE2* gene was identified using ChIP-seq data from IFNβ treated K562 cells^8^.

To identify candidate regulatory elements controlling the extended *ACE2* locus, including *ACE2* and *dACE2*, in primary airway cells, we dug deeper and conducted ChIP-seq for the chromatin marks H3K27ac (activate loci), H3K4me1 (enhancers), H3K4me3 (promoters) and Polymerase II (Pol II) loading, both in the absence and presence of IFNβ (Figure 1c Supplementary Figure 2). DNase I hypersensitive (DHS) sites from lung tissues^6^ and SAECs^7^ served as *bona fide* predictors of regulatory regions. In addition to *ACE2*, the neighboring *TMEM27* gene, an *ACE2* homologue, as well as *BMX* are activated by interferons (Supplementary Figure 2) suggesting the possibility of jointly used regulatory elements. *ACE2* and *TMEM27* originated from a gene duplication and their response to interferon is equivalent. The positions of the chromatin boundary factor CTCF suggests that *ACE2* and *TMEM27* are located within a sub-TAD (Supplementary Figure 2).

In agreement with earlier studies^4^, we identified DHS sites at ex1c and in intron 17 (Figure 1c). In addition, we identified DHS sites at ex1a, likely marking the genuine *ACE2* promoter, a distal site marking a possible enhancer and in intron 15 coinciding with activate marks. The DHS site in intron 9 overlaps with strong H3K4me3 marks, identifying it as a genuine promoter region (Figure 1c). This site is also decorated with H3K4me1 and H3K27ac marks and extensive Pol II loading, hallmarks of a complex promoter/enhancer. H3K27ac and Pol II loading was further induced by IFNβ, reflecting increased *ACE2* expression. IFNβ activates the transcription factors STATs 1, 2 and 5 and ChIP-seq experiments from K562 erythroid cells stimulated with IFNβ^8^ revealed preferential binding of STAT5 to the intronic promoter/enhancer (Figure 1d) further supporting regulation through the JAK/STAT pathway. The STAT1 locus served as a control for the binding of STAT transcription factors (Supplementary Figure 3). While the *ACE2* promoter associated with ex1a is marked by a DHS site and H3K4me1 marks, there is little evidence of H3K4me3 and H3K27ac marks. However, it is well known that there is no direct relationship between gene activity and the presence of these marks.

Based on the presence of DHS sites, activating chromatin marks and Pol II loading, either in combination or by themselves, we predict additional enhancers. A prominent candidate enhancer, marked by a DHS, H3K4me1 and Pol II loading is positioned approximately 16 kb distal of ex1a and additional putative regulatory regions are located with introns (Figure 1c). Activating histone marks and DHS sites^6,7^ are also present in the genuine *ACE2* promoter and the distal region in lung tissue (Supplementary Figure 4).

Interferons activate genetic programs through the JAK/STAT signaling pathway and JAK inhibitors are used clinically in COVID-19 patients in an effort to suppress the genomic consequences of cytokine storms^9,10,11^. To investigate if interferon-induced *ACE2* expression is controlled by the JAK/STAT pathway, we cultured SAECs in the presence of IFNβ and the JAK inhibitors, Baricitinib and Ruxolitinib, followed by RNA-seq and qRT-PCR assays (Supplementary Tables 9-10 and Figure 2) and ChIP-seq analyses (Figure 2). Both inhibitors suppressed the IFNβ-induced increase of the full-length *ACE2* (ex1a, ex1b and ex9) and the *dACE2* (ex1c) transcripts (Figure 2a and b), supporting that their respective promoters are under JAK/STAT control. The efficacy of the two inhibitors extended to a range of genetic programs activated through the pan JAK/STAT pathway (Figure 2c and Supplementary Figure 5) and induction of *bona fide* interferon stimulated genes (ISG), such as *ISG15*, was suppressed (Figure 2d). In *ACE2*, as in other ISGs, Ruxolitinib treatment mitigated the establishment of activating H3K27ac marks and Pol II loading over distal and intronic regulatory elements (Figure 2e-h).

**Figure 2.**
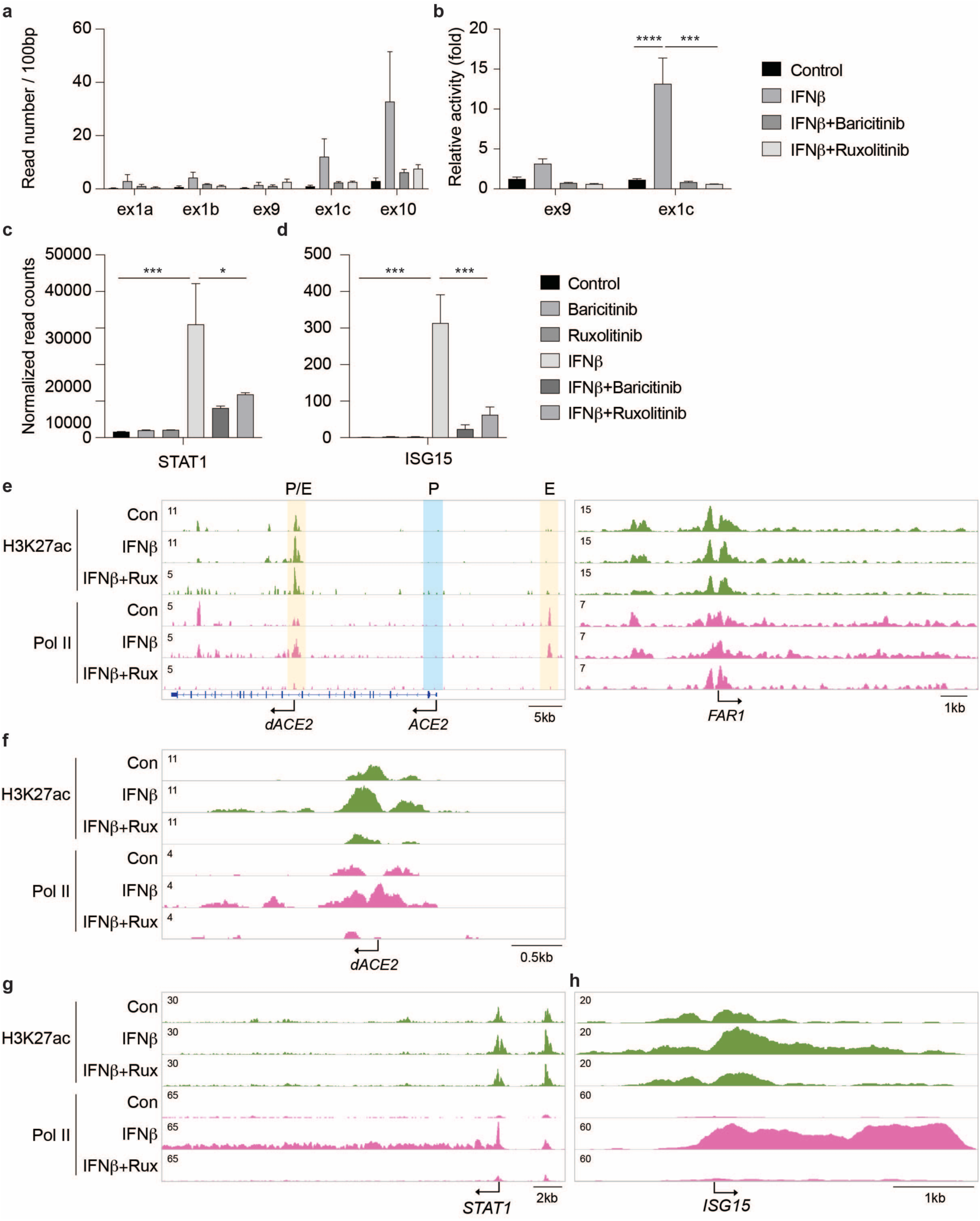
JAK inhibitors mitigated activation of IFNβ-stimulated genes. **a-b**. SAECs were cultured in the presence of IFNβ and JAK inhibitors, either Baricitinib or Ruxolitinib followed by RNA-seq analyses and qRT-PCR. Reads covering key exons are displayed. qRT-PCR results are shown as the means ± s.e.m. of independent biological replicates (Control and IFNβ, *n* = 9; IFNβ+JAK inhibitors, *n* = 3). Two-way ANOVA with followed by Tukey’s multiple comparisons test was used to evaluate the statistical significance of differences. **c-d**. *STAT1* and *ISG15* mRNA levels from control and experimental cells were measured by RNA-seq. Results are shown as the means ± s.e.m. of independent biological replicates (*n* = 3). One-way ANOVA with followed by Dunnett’s multiple comparisons test was used to evaluate the statistical significance of differences. **e-h**. H3K27ac marks, and Pol II loading at the *ACE2, STAT1* and *ISG15* loci in SAECs in the absence and presence of IFNβ and the JAK inhibitor, Ruxolitinib.

## Discussion

In summary, we have assessed the extended *ACE2* locus for regulatory elements controlling gene expression induced by interferons and other, not yet defined, stimuli. We show that the intronic regulatory region bears promoter and enhancer marks and binds STAT transcription factors, thus constituting a complex regulatory element controlling interferon-induced *dACE2* expression^4^. We identified additional candidate regulatory elements throughout the locus that can be invoked in controlling transcription of the full-length transcripts yielding the biologically active ACE2. Regulation of the human and mouse *ACE2*^5^ loci displays distinct differences, yet they share their response to cytokines and the JAK/STAT pathway and the mouse *Ace2* gene is activated in mammary tissue by cytokines through a JAK2/STAT5-dependent intronic enhancer^5^. Further studies in tissues and primary cells are needed to understand the complex, possibly cell-specific, regulation of the human *ACE2* locus in organs with extrapulmonary manifestations of SARS-CoV-2 infection. Our demonstration that JAK inhibitors mitigate interferon-induced activation of inflammatory programs has practical implications for their use in treating COVID-19 patients.

## Supplementary Information

### Methods

#### Cell culture

Human small airway epithelial cells (SAEC) obtained from Lifeline Technology (FC-0016) were expanded using the complete BronchiaLife^™^ media kit (Lifeline Technology, LL-0023). All culture wares were pre-coated in 30 *μ*g/ml of Fibronectin (ThermoFisher Scientific, 33016015) for at least 1h at room temperature. Calu-3 line (ATCC, HTB-55^™^) were cultured using Eagle’s Minimum Essential Medium (ATCC, 30-2003^™^) containing 10% fetal bovine serum (Cytiva, SH3007103) in 5% CO2 atmosphere at 37°C.

Cytokines (10 ng/ml; Human IFNβ, 300-02BC; Human IFNγ, 300-02; Human IL6, 200-06; Human IL7, 200-07; Human Growth hormone, 100-40, Peprotech; Human IFN α2b, 78077.1, Stem Cell Technologies; Human IFNλ3, 5259-IL-025, R&D systems) were treated in respective culture media and the cells were incubated for 12hr in 5% CO2 atmosphere at 37°C. The cells were washed with PBS (Gibco, 14190144) twice and harvested.

Jak inhibitors, 10 *μ*M of either Baricitinib (HY-15315A, MedChemExpress) or Ruxolitinib (HY-50856A, MedChemExpress), were added to BronchiLiafe ^™^ media with or without IFNβ. SAEC and incubated for 12hrs and then washed with PBS twice and harvested.

### RNA isolation and quantitative real-time PCR (qRT–PCR)

Total RNA was extracted from the collected cells and purified using the PureLink RNA Mini Kit (Invitrogen) according to the manufacturer’s instructions. cDNA was synthesized from total RNA using Superscript II (Invitrogen). Quantitative real-time PCR (qRT-PCR) was performed using TaqMan probes (ACE2, Hs01085333_m1; *STAT1*, Hs01013996_m1; *GAPDH*, Hs02786624_g1, Thermo Fisher scientific) on the CFX384 Real-Time PCR Detection System (Bio-Rad) according to the manufacturer’s instructions. Exon9 and exon1c mRNA were measured with the following primers, that were used by Onabajo et al.^4^, using SYBR Green system: Forward, 5’-GGGCGACTTCAGGATCCTTAT-3’, Reverse, 5’-GGATATGCCCCATCTCATGATGG-3’; Forward, 5’-GGAAGCAGGCTGGGACAAA-3’, Reverse, 5’-AGCTGTCAGGAAGTCGTCCATTG-3’. PCR conditions were 95°C for 30s, 95°C for 15s, and 60°C for 30s for 40 cycles. All reactions were done in triplicate and normalized to the housekeeping gene *GAPDH*. Relative differences in PCR results were calculated using the comparative cycle threshold (*C*_*T*_) method.

### Total RNA sequencing (Total RNA-seq) and data analysis

Total RNA was extracted from the collected cells and purified using the PureLink RNA Mini Kit (Invitrogen) according to the manufacturer’s instructions. Ribosomal RNA was removed from 1 μg of total RNAs and cDNA was synthesized using SuperScript III (Invitrogen). Libraries for sequencing were prepared according to the manufacturer’s instructions with TruSeq Stranded Total RNA Library Prep Kit with Ribo-Zero Gold (Illumina, RS-122-2301) and paired-end sequencing was done with a HiSeq 3000 instrument (Illumina).

Total RNA-seq read quality control was done using Trimmomatic^12^ (version 0.36) and STAR RNA-seq^13^ (version STAR 2.5.4a) using 50bp paired-end mode was used to align the reads (hg19). HTSeq^14^ was to retrieve the raw counts and subsequently, R (https://www.R-project.org/), Bioconductor^15^ and DESeq2^16^ were used. Additionally, the RUVSeq^17^ package was applied to remove confounding factors. The data were pre-filtered keeping only those genes, which have at least ten reads in total. Genes were categorized as significantly differentially expressed with an adjusted p-value (pAdj) below 0.05 and a fold change > 2 for up-regulated genes and a fold change of < −2 for down-regulated ones. The visualization was done using dplyr (https://CRAN.R-project.org/package=dplyr) and ggplot2^18^. Sequence read numbers were calculated using Samtools^19^ software with sorted bam files.

### Chromatin immunoprecipitation sequencing (ChIP-seq) and data analysis

Chromatin was fixed with formaldehyde (1% final concentration) for 15 min at room temperature, and then quenched with glycine (0.125 M final concentration). Samples were processed as previously described^20^. The following antibodies were used for ChIP-seq: H3K27ac (Abcam, ab4729), RNA polymerase II (Abcam, ab5408), H3K4me1 (Active Motif, 39297) and H3K4me3 (Millipore, 07-473). Libraries for next-generation sequencing were prepared and sequenced with a HiSeq 3000 instrument (Illumina). Quality filtering and alignment of the raw reads was done using Trimmomatic^12^ (version 0.36) and Bowtie^21^ (version 1.1.2), with the parameter ‘-m 1’ to keep only uniquely mapped reads, using the reference genome hg19. Picard tools (Broad Institute. Picard, http://broadinstitute.github.io/picard/. 2016) was used to remove duplicates and subsequently, Homer^22^ (version 4.8.2) software was applied to generate bedGraph files. Integrative Genomics Viewer^23^ (version 2.3.81) was used for visualization.

## Statistical anlaysis

For comparison of samples, data were presented as standard deviation in each group and were evaluated with a two-way ANOVA followed by Tukey’s multiple comparisons test or a one-way ANOVA with Dunnett’s multiple comparisons test using PRISM 8 GraphPad (version 8.2.0). Statistical significance was obtained by comparing the measures from wild-type or control group, and each mutant group. A value of \**P* < 0.05, \*\**P* < 0.001, \*\*\**P* < 0.0001, \*\*\*\**P* < 0.00001 was considered statistically significant.

## Data availability

All data were obtained or uploaded to Gene Expression Omnibus (GEO). ChIP-seq for STATs was obtained under GSE31477. ChIP-seq data for H3K27ac and H3K4me1 from human lung tissues were downloaded from GSE143115 and 142958. DNase I hypersensitive (DHS) data from human lung tissues and SAEC were obtained under GSE90364 and 29692, respectively. The RNA-seq and ChIP-seq data from SAEC will be uploaded in GEO before publishing the manuscript.

## Acknowledgments

We thank Ilhan Akan, Sijung Yun and Harold Smith from the NIDDK genomics core for NGS and NCATS Chemical Genomics Center Team for the JAK inhibitors. This work utilized the computational resources of the NIH HPC Biowulf cluster (http://hpc.nih.gov).

## Funding

This work was supported by the Intramural Research Programs (IRPs) of National Institute of Diabetes and Digestive and Kidney Diseases (NIDDK) and National Center for Advancing Translational Sciences (NCATS).

## Author contribution

HKL: project conception, experimental design and execution, data analysis, preparation of figures, writing manuscript; OJ: experimental design and execution; LH: project conception, experimental design, data analysis, preparation of figures, writing manuscript.

## Competing interests

The authors declare no competing financial interests.

---

**Supplementary Table 1**. mRNA levels of genes associated with the pan JAK-STAT pathway in SAEC.

**Supplementary Table 2**. List of all genes with normalized read counts in each replicate at Control and IFN α treated SAEC, log_2_ (fold change), *p*-value and adjusted *p*-value as well as upregulated gene list and GSEA analysis.

**Supplementary Table 3**. List of all genes with normalized read counts in each replicate at Control and IFNβ treated SAEC, log_2_ (fold change), *p*-value and adjusted *p*-value as well as upregulated gene list and GSEA analysis.

**Supplementary Table 4**. List of all genes with normalized read counts in each replicate at Control and IFNγ treated SAEC, log_2_ (fold change), *p*-value and adjusted *p*-value as well as upregulated gene list and GSEA analysis.

**Supplementary Table 5**. List of all genes with normalized read counts in each replicate at Control and IFNλ3 treated SAEC, log_2_ (fold change), *p*-value and adjusted *p*-value as well as upregulated gene list and GSEA analysis.

**Supplementary Table 6**. List of all genes with normalized read counts in each replicate at Control and IL6 treated SAEC, log_2_ (fold change), *p*-value and adjusted *p*-value as well as upregulated gene list and GSEA analysis.

**Supplementary Table 7**. List of all genes with normalized read counts in each replicate at Control and IL7 treated SAEC, log_2_ (fold change), *p*-value and adjusted *p*-value as well as upregulated gene list and GSEA analysis.

**Supplementary Table 8**. List of all genes with normalized read counts in each replicate at Control and GH treated SAEC, log_2_ (fold change), *p*-value and adjusted *p*-value as well as upregulated gene list and GSEA analysis.

**Supplementary Table 9**. List of all genes with normalized read counts in each replicate at IFNβ and Baricitinib with IFNβ, treated SAEC, log_2_ (fold change), *p*-value and adjusted *p*-value as well as upregulated gene list and GSEA analysis.

**Supplementary Table 10**. List of all genes with normalized read counts in each replicate at IFNβ and Ruxolitinib with IFNβ, treated SAEC, log_2_ (fold change), *p*-value and adjusted *p*-value as well as upregulated gene list and GSEA analysis.

**Supplementary Figure 1.**
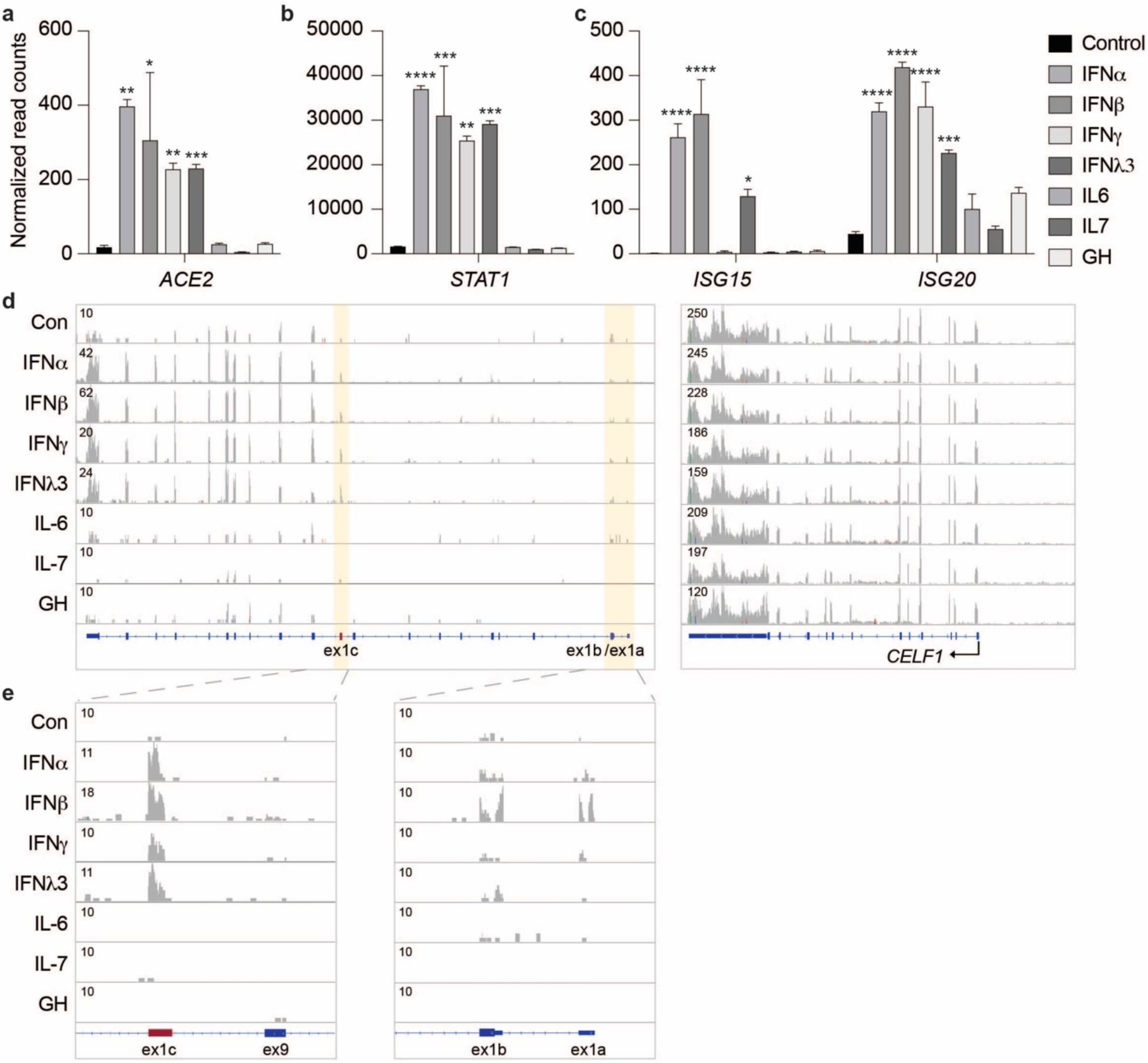
Induction of *ACE2* expression by interferons. **a-c**. *ACE2*, and *STAT1* and *ISGs* mRNA levels from control cells and cells treated with different cytokines were measured by RNA-seq. Results are shown as the means ± s.e.m. of independent biological replicates (*n* = 3). One-way ANOVA followed by Dunnett’s multiple comparisons test was used to evaluate the statistical significance of differences. **d-e**. RNA-seq reads are matched with exons.

**Supplementary Figure 2.**
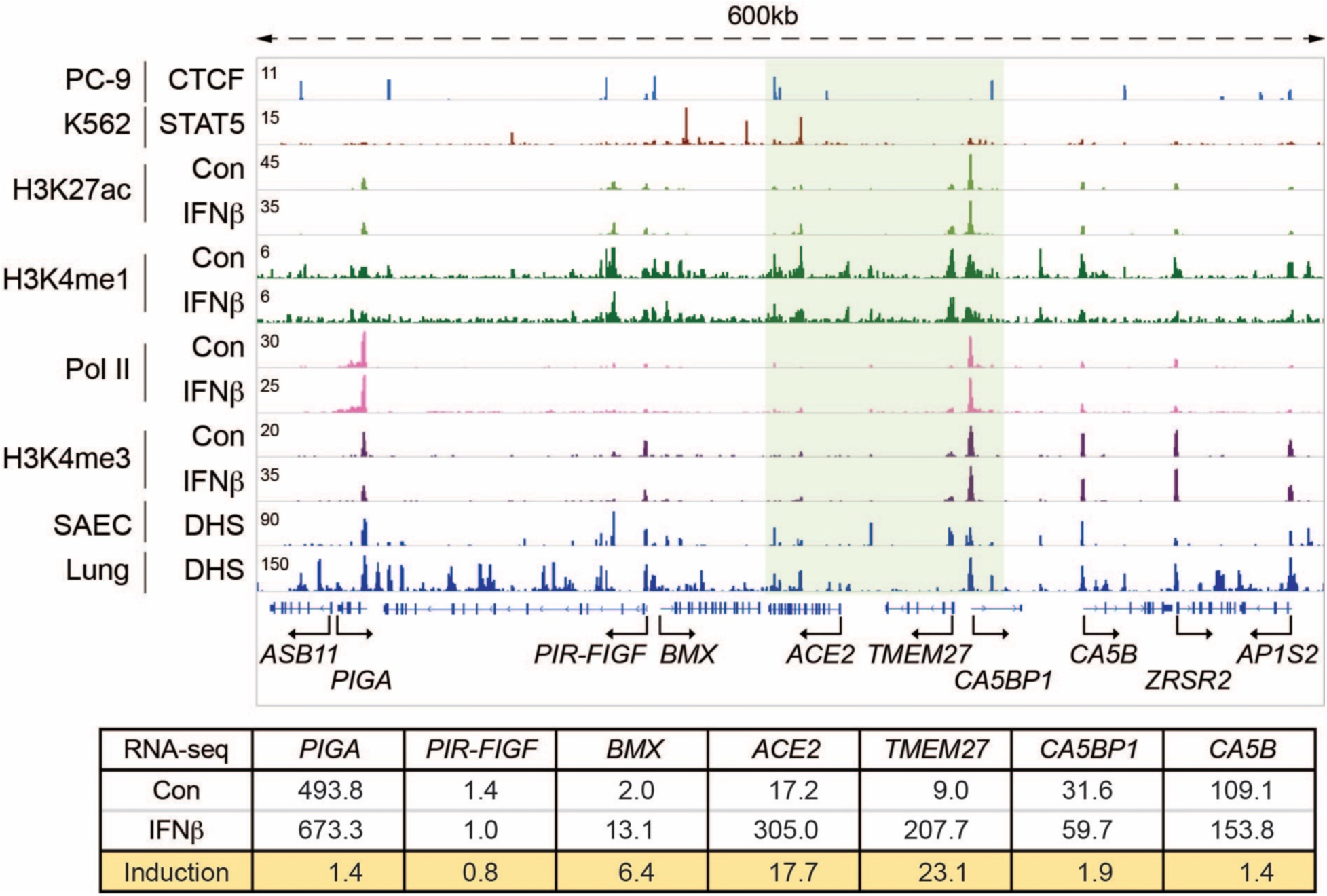
Structure and expression of the extended ACE2 locus in primary airway cells. Regulatory marks in the 600 kb locus including *ACE2* and neighboring genes in SAECs. mRNA levels in the absence and presence of IFNβ were measured by RNA-seq. ChIP-seq for H3K4me1, H3K27ac, H3K4me3 and Pol II was conducted in SAECs in the absence and presence of IFNβ. CTCF and STAT5 ChIP-seq data and DHS data are from ENCODE.

**Supplementary Figure 3.**
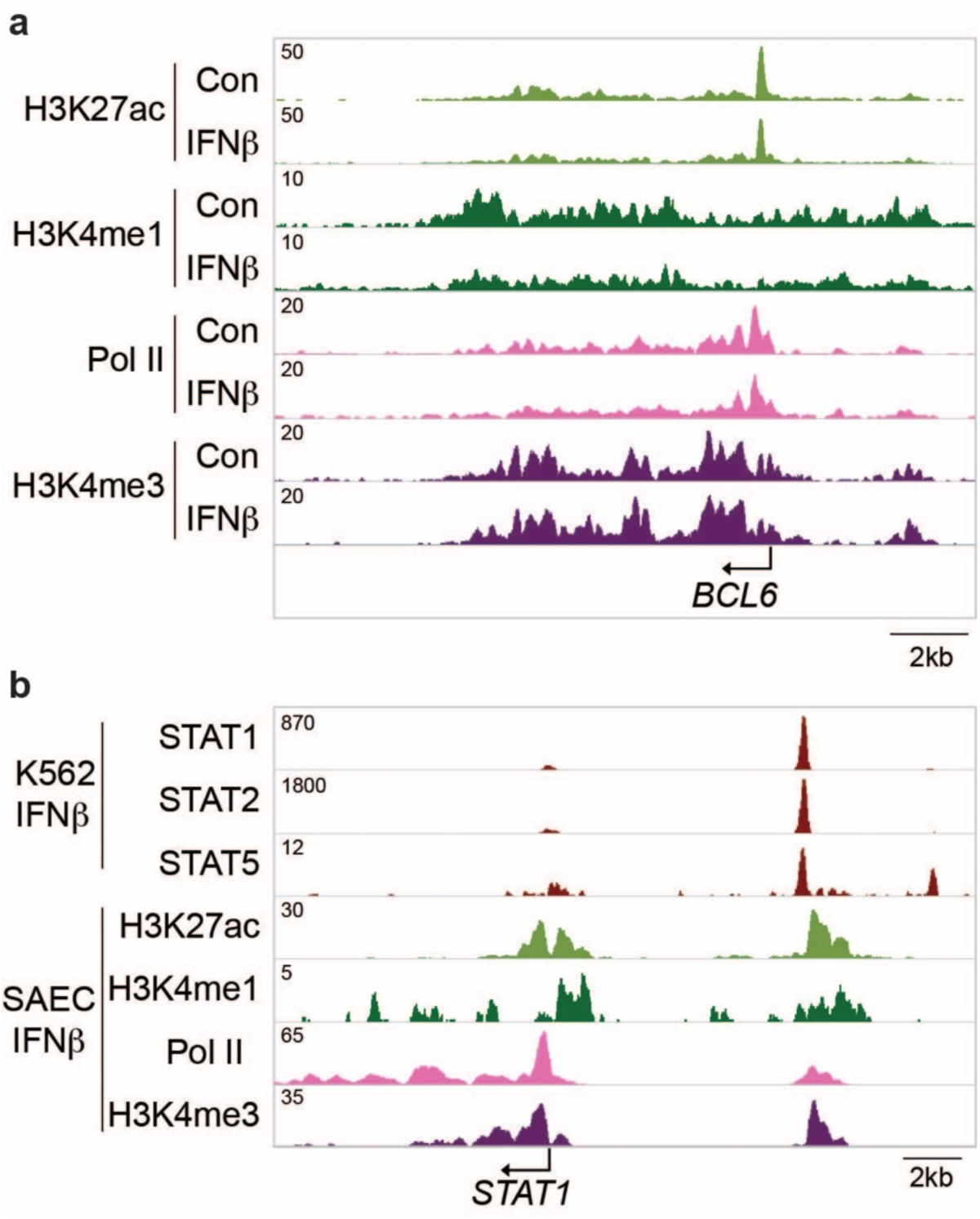
Activating histone marks and Pol II loading at control loci. **a**. The *BCL6* severed as ChIP-seq control for histone marks and RNA Pol II. Solid arrows indicate the orientation of genes. **b**. ChIP-seq data for STAT transcription factors, histone marks and RNA Pol II on the *STAT1* locus.

**Supplementary Figure 4.**
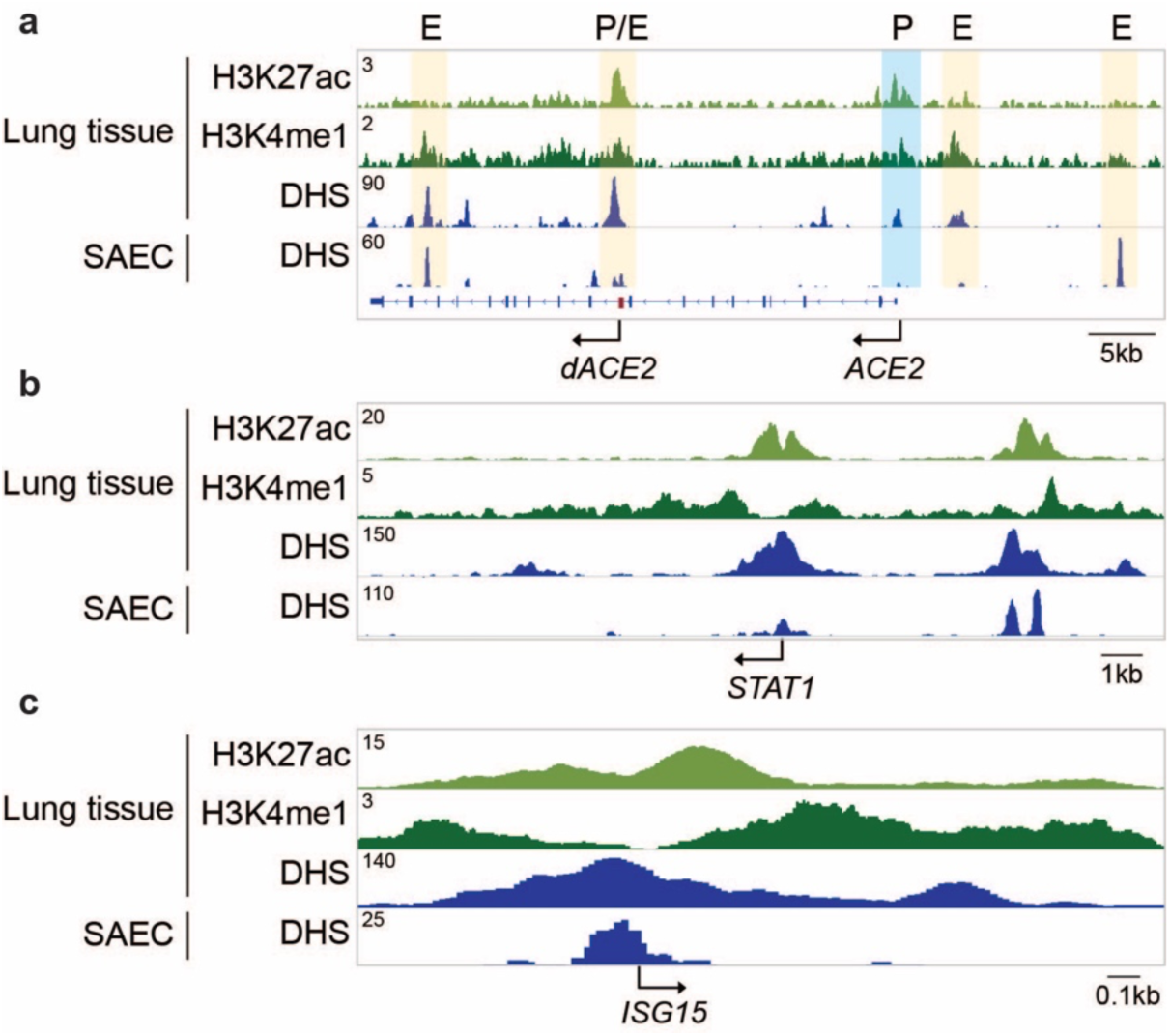
Activating histone marks at the *ACE2* locus in lung tissue. ChIP-seq and DHS from human lung tissues and DHS from SAECs display regulatory elements at the *ACE2, STAT1* and *ISG15* loci.

**Supplementary Figure 5.**
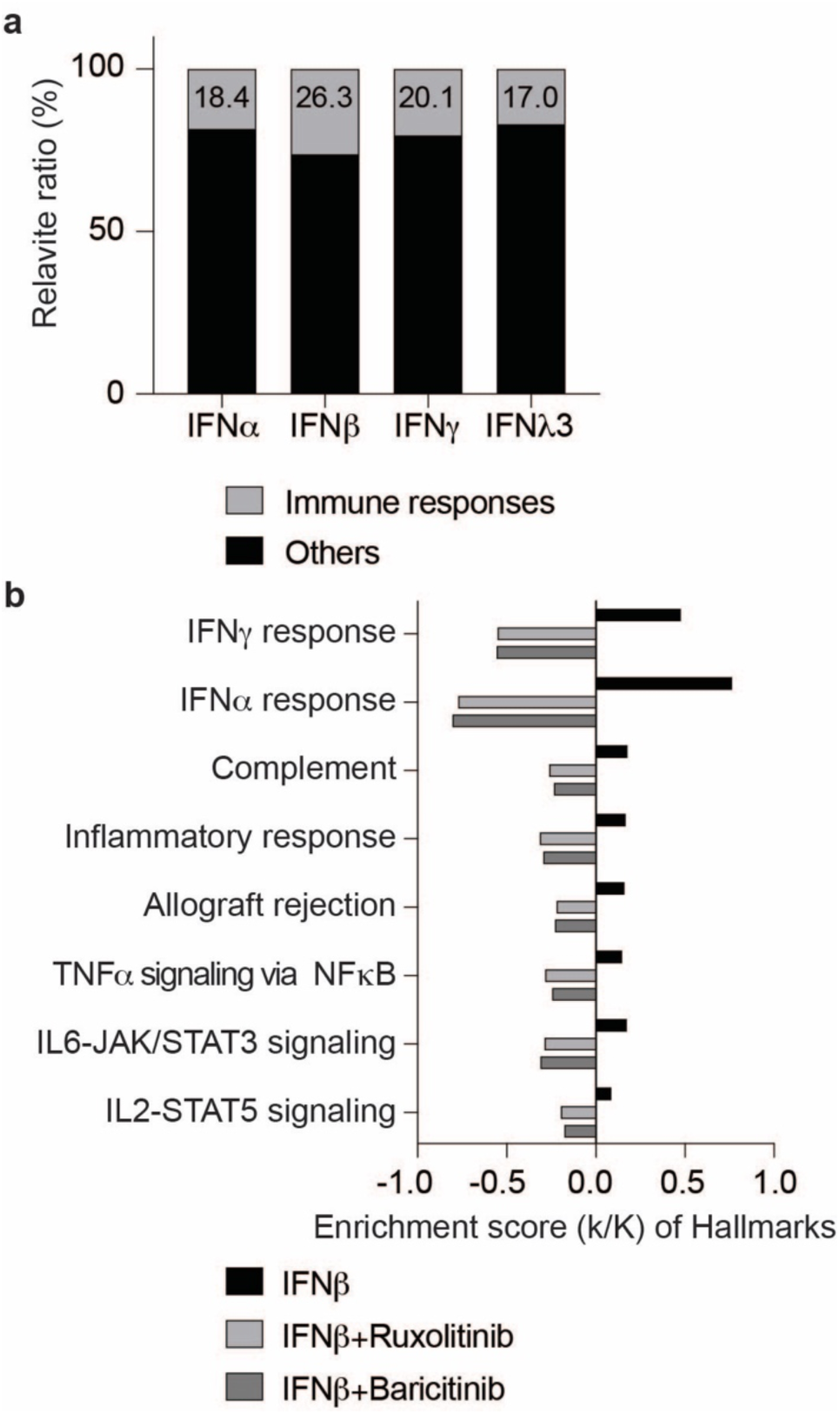
Activation of immune response genes by Interferons in human primary airway epithelium (SAEC) is mitigated by JAK inhibitors. **a**. Genes induced significantly by IFN α, β, γ and λ3 were significantly enriched in Hallmark Gene Sets (FDR q-value < 0.005). Immune response genes accounted for 18.4%, 26.3%, 20.1% and 17% of the upregulated genes. **b**. Expression of immune pathway genes induced by IFNβ was mitigated by the JAK inhibitors Ruxolitinib and Baricitinib.

## Notes

### Competing Interest Statement

The authors have declared no competing interest.

